# Cold tolerance of mountain stoneflies (Plecoptera: Nemouridae) from the high Rocky Mountains

**DOI:** 10.1101/2020.06.25.171934

**Authors:** Scott Hotaling, Alisha A. Shah, Michael E. Dillon, J. Joseph Giersch, Lusha M. Tronstad, Debra S. Finn, H. Arthur Woods, Joanna L. Kelley

## Abstract

How aquatic insects cope with cold temperatures is poorly understood. This is particularly true for high-elevation species that often experience a seasonal risk of freezing. In the Rocky Mountains, nemourid stoneflies (Plecoptera: Nemouridae) are a major component of mountain stream biodiversity and are typically found in streams fed by glaciers and snowfields, which due to climate change, are rapidly receding. Predicting the effects of climate change on mountain stoneflies is difficult because their thermal physiology is largely unknown. We investigated cold tolerance of several alpine stoneflies (*Lednia tumana, Lednia tetonica*, and *Zapada* spp.) from the Rocky Mountains, USA. We measured the supercooling point (SCP) and tolerance to ice enclosure of late-instar nymphs collected from a range of thermal regimes. SCPs varied among species and populations, with the lowest SCP measured for nymphs from an alpine pond, which is much more likely to freeze solid in winter than flowing streams. We also show that *L. tumana* cannot survive being enclosed in ice, even for short periods of time (less than three hours) at relatively mild temperatures (−0.5 °C). Our results indicate that high-elevation stoneflies at greater risk of freezing may have correspondingly lower SCPs, and despite their common association with glacial meltwater, they appear to be living near their lower thermal limits.

## INTRODUCTION

Freshwater habitats in cold regions typically experience winter ice cover and freezing temperatures that can be injurious or lethal to aquatic insects (Danks 2008). Resident species may avoid extreme cold by moving to warmer or more thermally stable microclimates or by modifying local conditions (e.g., through case-making; Danks 1971). Others withstand cold conditions *in situ* through supercooling—maintaining internal water in liquid form below its freezing point—or freeze tolerance (Danks 2008, Sinclair et al. 2015, Zachariassen 1985). Organisms living in wet environments are likely to encounter external ice when their body temperatures are already at or near freezing. Thus, inoculative freezing—in which contact with external ice induces the formation of internal ice—is likely the main driver of organismal freezing in these habitats (Frisbie & Lee Jr. 1997).

At high elevations, aquatic habitats are typically distributed across a mosaic of streams and ponds that are fed by an array of hydrological sources (e.g., glaciers, perennial snowfields, subterranean ice; Hotaling et al. 2017) and remain snow-covered for much of the year (Lencioni 2004). Mountain streams appear to not freeze solid because of their steep gradients and year-round flow, with minimum temperatures often close to 0 °C in the main channel (Giersch et al. 2017, Shah et al. 2017, Tronstad et al. in press). Resident taxa are therefore predicted to experience minimum temperatures of ∼0 °C. This prediction rests on the assumption that overwintering nymphs remain in the unfrozen primary stream channel throughout the winter season. However, because of the inherent variation in aquatic habitats, nymphs inhabiting different regions of a stream (e.g., littoral zone, bed sediments) may experience subfreezing temperatures, even when the rest of the stream remains unfrozen (Lencioni 2004). Alpine ponds, in contrast, have little or no flow and are often shallow, making them susceptible to freezing solid (Wissinger et al. 2016). As melt-induced flows are reduced in the future under climate change (Huss & Hock 2018), headwater streams will become shallower with less flow velocity, raising the risk of freezing in all seasons for resident organisms.

Stoneflies (Order Plecoptera) inhabit every continent except Antarctica and, when present, often represent a substantial portion of aquatic biodiversity (DeWalt et al. 2015). After hatching, stoneflies follow a two-stage life cycle with development through multiple instars as aquatic, larval nymphs followed by emergence as winged, terrestrial adults. Whether development occurs in a single year or spans multiple seasons, eggs and larvae of high-elevation stoneflies are exposed to near freezing temperatures. Cold tolerance of stoneflies and of aquatic insects broadly is poorly known, particularly for non-adult life stages (i.e., eggs or nymphs; Danks 2008) and species living at high latitudes and elevations—where taxa are most likely to experience freezing. Aquatic insects generally employ an array of physiological and biochemical strategies to mitigate cold stress (reviewed by Lencioni et al. 2004). These include the production of cryoprotectants to lower the temperature of internal ice formation (Duman et al. 2015), cryoprotective dehydration where body water is lost to lower the amount available for freezing (e.g., Gehrken & Sømme 1987), and the ability to resist anoxic conditions induced by ice enclosure (e.g., Brittain & Nagell 1981). Eggs of the stonefly *Arcynopteryx compacta* (Perlodidae) from the mountains of southern Norway avoid freezing by supercooling to -31 °C, a trait mediated by the loss of approximately two-thirds of the eggs’ water content (Gehrken 1989, Gehrken & Sømme 1987). Nymphs of another high-latitude stonefly, *Nemoura arctica* (Nemouridae), survive freezing to well below 0 °C by preventing intracellular ice formation through the production of a xylomannan-based glycolipid, which inhibits inoculative freezing through the inactivation of ice nucleating agents (Walters et al. 2009, 2011). However, it is unlikely that glycolipid production represents a singular ‘magic bullet’ that confers freeze tolerance in stoneflies, or aquatic insects broadly, given the diversity of known freeze protecting molecules (Toxopeus & Sinclair 2018). Like *N. arctica*, high-alpine nemourids in North America appear to overwinter as early-instar nymphs (Figure S1) and may be exposed to a similar risk of freezing. How they cope with such risks, however, has not been investigated.

In this study, we investigated cold tolerance across populations of late-instar nymphs from at least three species [*Lednia tumana* (Ricker 1952), *Lednia tetonica* Baumann & Call 2010, and *Zapada* spp.; Nemouridae)] in the northern Rocky Mountains, USA. For all populations, we measured the dry supercooling point (SCP)—the temperature at which an organism releases latent heat indicative of ice formation (Sinclair et al. 2015)—in the absence of ice. Enclosure in ice may also be a major risk for mountain stoneflies, especially in slower flowing shallow streams. Ice-enclosure can cause mortality through inoculative freezing, hypoxia, mechanical damage, or a combination of factors (Conradi-Larsen & Sømme 1973). One nemourid, *N. arctica*, can survive ice enclosure (Walters et al. 2009), but how widespread this ability is among stoneflies is unknown. Thus, for one population from Lunch Creek, we also tested whether nymphs could survive ice enclosure. Given the perennial and fast-flowing conditions of alpine streams, we expected winter conditions to be largely constant with temperatures near 0 °C and likelihood of freezing to generally be low. Therefore, we did not expect stonefly SCPs to vary among stream-inhabiting populations. Conversely, because high elevation pond-dwelling stoneflies likely experience a greater risk of freezing, we expected resident nymphs to exhibit lower SCPs. Furthermore, given that *N. arctica* nymphs can survive being enclosed in ice and that streams containing *Lednia* are commonly near 0 °C even in summer (Figure S2), we expected *L. tumana* nymphs to survive ice enclosure. Our study represents a preliminary step toward understanding how alpine stonefly cold tolerance varies at broad taxonomic and environmental scales—i.e., across species, populations, and habitats— setting the stage for future, more targeted studies of the group.

## MATERIALS AND METHODS

### Sampling and environmental data collection

For SCP measurements, we collected late-instar stonefly nymphs from six populations in the Rocky Mountains between 29 July – 6 August 2018 (Figure 1; Tables 1, S1). To test whether high-elevation nemourids could survive being enclosed in ice, we collected additional late-instar *L. tumana* nymphs from Lunch Creek in Glacier National Park, MT, on 17 August 2019. Nymphs were transferred in 1 L Whirl-Pak bags (Nasco) filled with streamwater from the backcountry to the laboratory in ice-filled coolers. Transport duration depended on distance from a trailhead, ranging from a few hours (Lunch Creek) to more than 24 hours (Mount Saint John). Populations were sampled from two Rocky Mountain sub-ranges centered around Glacier and Grand Teton National Parks (Figure 1). Due to the remote, rugged terrain and high seasonal snowfall in the Rocky Mountains, we targeted our specimen collections to occur in late summer, soon after most streams in our study area became free of seasonal snow. We identified specimens based on morphology and collection localities following previous studies (e.g., Giersch et al. 2017). Unlike *Lednia*, multiple *Zapada* species can be present in the same stream, and therefore, we cannot exclude the presence of more than one species in the Wind Cave population (Table 1).

**Table 1.** Environmental variation and habitat types included in this study. T_MIN_ and T_MEAN_: the minimum and mean temperatures observed for each site, respectively. C: specific conductivity (*μ*S cm^-1^); PI: Pfankuch Index, a measure of stream channel stability (higher values correspond to a less stable streambed); SCP: mean and standard deviation of the supercooling point. The focal 24-hour period for temperature measurement was 31 July 2019 for all sites except Lunch Creek (31 July 2014) and Wind Cave (28 July 2019). All temperatures (T_MIN_, T_MEAN_, SCP) are in degrees Celsius. All sites are in Grand Teton National Park except Lunch Creek (Glacier National Park).

| Population | Taxon | Type | $T_{\text{MIN}}$ | $T_{\text{MEAN}}$ | C | PI | SCP |
| --- | --- | --- | --- | --- | --- | --- | --- |
| Lunch Creek (LC) | *L. tumana* | Snowmelt | 4.2 | 6.2 | 40.8 | 25 | $-5.9 \pm 2.0$ |
| Wind Cave (WC) | *Zapada* spp. | Icy seep | 2.6 | 2.8 | 101.1 | 18 | $-7.2 \pm 2.0$ |
| Mt. Saint John (MSJ) | *L. tetonica* | Icy seep | 2.4 | 3.0 | 25.0 | 34 | $-5.8 \pm 0.7$ |
| Cloudveil Dome (CD) | *L. tetonica* | Glacier-fed | 1.8 | 2.0 | 4.1 | 32 | $-7.0 \pm 0.9$ |
| Skillet Glacier (SG) | *L. tetonica* | Glacier-fed | 2.7 | 4.4 | 3.1 | 34 | $-5.2 \pm 1.1$ |
| Tetonica Pond<sup>a</sup> (TP) | *L. tetonica* | Pond | 2.6 | 3.1 | 29.3 | n/a | $-7.5 \pm 1.3$ |

<sup>a</sup> Named by the authors. Does not reflect official conventions.

**Figure 1.**
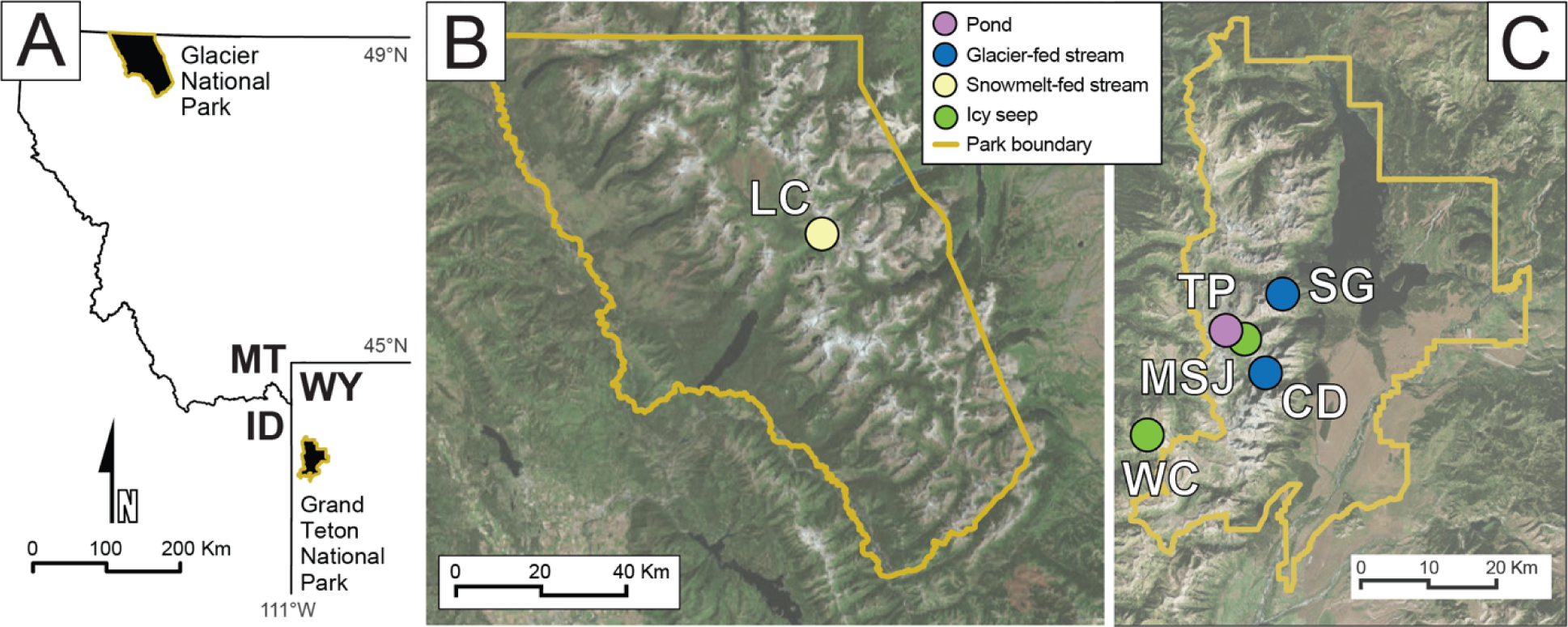
Populations and habitat types included in this study. A) Locations of the focal ranges in the Rocky Mountains, B) Glacier National Park, Montana, USA, and C) Grand Teton National Park and surrounding mountains, Wyoming, USA. See Table 1 for complete population names.

We collected environmental data at each site to characterize the temperatures nymphs experience in the wild and to categorize sites by habitat type (e.g., glacier-fed, snowmelt-fed, groundwater-fed, or a high-elevation pond; see below). We recorded temperature hourly with HOBO loggers (Onset Computer Corporation) placed within ∼5 cm of the main channel streambed. Lengths of logger deployments ranged from less than 24 hours (Mount Saint John, Tetonica Pond) to a full calendar year (Lunch Creek, Skillet Glacier, Wind Cave). Using these data, we constructed 24-hour thermographs for each site on a representative day in late July. Our focal 24-hour period was 31 July 2019 for all sites except Lunch Creek (31 July 2014) and Wind Cave (28 July 2019). For the focal period, we measured the lowest (T_MIN_) and mean (T_MEAN_) temperatures. We also measured specific conductivity (C) with a YSI Professional ProPlus multiparameter probe, which was calibrated at the trailhead before each trip. Specific conductivity reflects ionic concentration in the water and is elevated in water that interacts with inorganic debris (e.g., streams fed by debris-covered rock glaciers). We assessed stream channel stability via a modified version of the Pfankuch Index (PI, Peckarsky et al. 2014). We estimated when sites became ice-free in 2018 using satellite imagery (Copernicus Sentinel 2).

We classified sites into one of four habitat types based on a combination of primary hydrological source, environmental variation, geomorphology, and field surveys. Habitat types included: streams fed by a surface glacier (e.g., “glacier-fed”), streams fed by a perennial snowfield (“snowmelt-fed”), streams emanating from subterranean ice (e.g., rock glaciers, “icy seep”; Hotaling et al. 2019, Tronstad et al. in press), and slow-flowing, alpine ponds (“pond”), identified by their low-angle profile and appearance of standing water. To discern among the three stream types, we relied on geographic data paired with the environmental signature of primary hydrological sources—glaciers, snowfields, or subterranean ice masses. Glacier-fed streams were identified by four criteria: named glaciers feeding them, low minimum temperatures (T_MIN_ < 3 °C), low C (< 10 *μ*S cm^-1^), and high PI (> 25, indicating more unstable stream channels). Streams were classified as snowmelt-fed if a permanent snowfield was visibly feeding them, they exhibited higher minimum temperatures (T_MIN_ > 3 °C), moderate C (10-50 *μ*S cm^-1^), and a moderately stable stream channel (PI = 18-25). We categorized streams as icy seeps if they were consistently low (e.g., T_MEAN_ ≤ 3 °C) yet had no named glaciers or significant perennial snowfields feeding them, a classic “lobe” of a rock glacier was visible (e.g., Mount Saint John), and they exhibited moderate to high C for mountain streams (> 25 *μ*S cm^-1^).

### Measuring supercooling points

In the laboratory, nymphs were held at 3 °C with no access to food for 12-72 hours (Table S1). For each SCP measurement, we blotted nymphs with Kimwipes (Kimberly Clark) and placed them in dry 1.5 mL microcentrifuge tubes. A 30-gauge T-type thermocouple (2 mm soldered tip) was held in contact with each nymph’s abdomen by a piece of cotton wedged into the tube. Thermocouples were attached to 4-channel loggers (UX-120, ± 0.6 °C accuracy, Onset Corporation). Up to 12 microcentrifuge tubes were placed in an aluminum block in contact with a thermoelectric cooler, attached to a heat sink immersed in a glycol bath on the opposite face. We used a custom temperature controller to decrease the temperature at ∼0.5-1 °C per minute. We estimated the SCP as the lowest recorded temperature before observing the transient increase in temperature as the body released latent heat and started to freeze. After SCPs were recorded for all nymphs in a given experimental group, we promptly removed them from the experimental bath. Nymphs were clearly dead upon removal from the microcentrifuge tubes after the SCP tests, with no movement for several minutes and no response to probing. We subsequently measured body length to the nearest millimeter for all specimens with a microscope and ocular scale.

### Testing tolerance to ice enclosure

We assessed whether late-instar *L. tumana* nymphs could survive being enclosed in ice, a potential environmental risk factor in high-elevation streams and ponds. After collection, nymphs were maintained at 5 °C in the laboratory for 72 hours with no access to food. We then placed a single nymph in each well of an ice-cube tray with approximately 22 mL of streamwater. Nymphs were assigned to one of three treatments: -0.5 °C (freezing, *N* = 8), 0.1 °C (near freezing, *N* = 13), or control (5 °C, *N* = 6). Thermocouples were inserted into the wells to monitor temperature. Once the streamwater reached the desired temperature, we began the treatment which lasted for three hours. Following the experiment, we placed nymphs at 5 °C to recover for seven hours. After the recovery period, we assessed survival by observation and gentle prodding.

### Statistical analyses

We performed all analyses in R (R Core Team 2017). We first conducted an ANOVA to assess the effects of body length on SCP across all species. Next, to test if there were differences in SCP across populations, we conducted an ANOVA using population as the predictor variable and SCP as the response variable. A Tukey HSD correction for multiple comparisons was used to compare the mean SCP for each pair of streams. We used the same approach (ANOVA with Tukey HSD correction) to estimate the effect of habitat type (glacier-fed, snowmelt-fed, icy seep, or pond) or species (*L. tumana, L. tetonica*, or *Zapada* spp.) on SCP. Finally, we tested for population-specific variation in SCP for *L. tetonica*. We included body length as a covariate in all models to account for body size effects.

## RESULTS

### Environmental variation and habitat types

The sites sampled in this study represented four habitat types (Table 1). Tetonica Pond was the only site that lacked steep, flowing water, and thus was the only ‘pond’ we included. We also sampled two glacier-fed streams (Cloudveil Dome, Skillet Glacier outlet), two icy seeps (Wind Cave, Mount Saint John), and one snowmelt-fed stream (Lunch Creek). Over the focal 24-hour period, minimum temperature was lowest at Cloudveil Dome (T_MIN_ = 1.8 °C) and highest at Lunch Creek (T_MIN_ = 4.2 °C; Table 1). Mean temperature followed the same pattern with Cloudveil Dome the coldest (T_MEAN_ = 2.0 °C) and Lunch Creek the warmest (T_MEAN_ = 6.2 °C; Table 1). For all sites except Wind Cave, seasonal snow cover persisted well into July (Table S1). For Lunch Creek, Wind Cave, and Skillet Glacier, full-year thermographs indicate that the main stream channel remains unfrozen year-round (Figure S2), but whether this lack of freezing extends to the streambed and margins is unknown.

### Supercooling points

Across all species, populations, and habitats, the mean SCP was -6.5 ± 1.7 °C (Table 1). Larger nymphs had higher SCPs than smaller nymphs (*F*_(1,101)_ = 7.14, *P* = 0.009; Figure S3); we therefore included body length as a covariate in our statistical models. We found significant differences in SCPs among species (*F* _(2,109)_ = 3.25, *P* = 0.043; Figure 2A), populations (*F* _(7,95)_ = 4.528, *P*, ANOVA < 0.001), and habitats (*F* _(4,98)_ = 4.556, *P* = 0.002). Within a single species, *L. tetonica*, the SCP varied significantly among habitats (*F* _(3,62)_ = 10.4, *P* < 0.001; Figure 2B). Individuals from Tetonica Pond had the lowest mean SCP overall (−7.5 ± 1.3°C, Table 1) and those from glacier-fed streams had the highest (Table 1).

**Figure 2.**
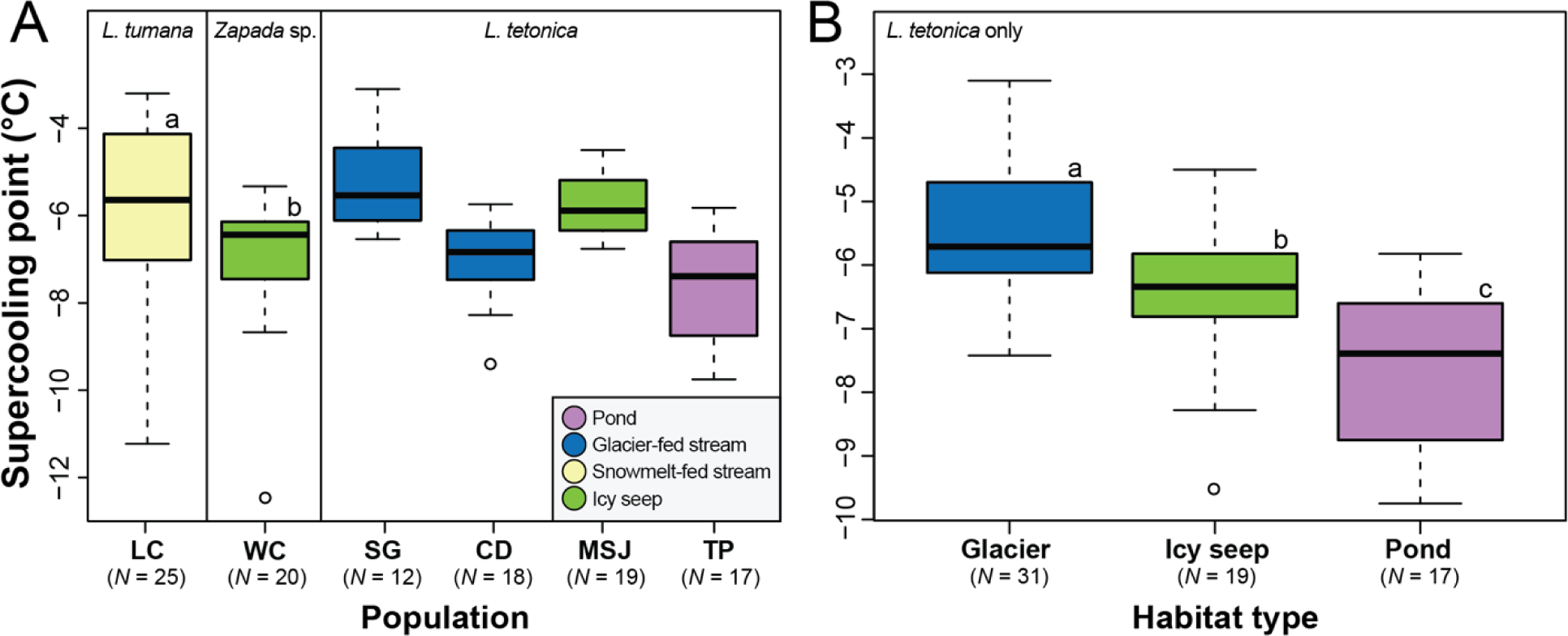
Box-and-whisker plots of the supercooling point (SCP) for alpine stonefly nymphs in the Rocky Mountains grouped by (A) species and population and (B) habitat type for *Lednia tetonica* only. In each plot, groups of lower-case letters are significantly different at Tukey’s *P* < 0.05 (see Table S2 for detailed statistics). Dark lines represent median values in the data. Upper and lower bounds of the box are the upper and lower quartiles, respectively. Outliers are indicated by open circles. Population acronyms: Lunch Creek (LC), Wind Cave (WC), Skillet Glacier (SG), Cloudveil Dome (CD), Mount Saint John (MSJ), and Tetonica Pond (TP).

### Tolerance to ice enclosure

No *L. tumana* nymphs (*N =* 0 of 8) in the freezing treatment (−0.5 °C) survived being entirely enclosed in ice. In the near-freezing treatment (0.1 °C), ∼84% of nymphs (*N* = 7 of 8) survived. A thin layer of ice formed on the surface of the well during the treatment but the remainder of the streamwater beneath it containing each nymph did not freeze. All nymphs survived the 5 °C control treatment (*N* = 6 of 6).

## DISCUSSION

Stoneflies are a major component of aquatic biodiversity in mountain ecosystems yet little is known of their ability to cope with low temperatures, particularly as it relates to potential freezing stress. In this study, we measured supercooling points (SCPs) and tolerance to ice enclosure for high-elevation stonefly nymphs in the Rocky Mountains to link their physiological traits to the thermal regimes they experience. Our SCP estimates aligned with those described for other aquatic insects (−3.3° to -7.4 °C; Moore & Lee Jr 1991) and varied by species, population, and habitat. Because SCPs are typically higher for aquatic insects in summer versus winter, the values observed in our study likely reflect maximum dry SCPs for our focal species (Moore & Lee Jr 1991). Within *L. tetonica*, the alpine pond population exhibited lower SCPs than those from icy seeps (and the lowest mean SCP of any population, –7.5 ± 1.3 °C), which in turn had lower SCPs than those from glacier-fed streams. Based on our results, we cannot disentangle the mechanism driving these patterns of cold tolerance (i.e. plasticity or local adaptation), but a correlation between thermal regime and SCP appears to exist. Although hydrologically connected to steep, fast-flowing streams, the small size, shallowness, and extremely slow flow of Tetonica Pond likely elevates winter freezing risk for resident taxa; alpine ponds in the Rocky Mountains often freeze solid in winter (Wissinger et al. 2016). Snow cover persisted on Tetonica Pond until mid-July in 2018 (Table S1), suggesting it also may have been frozen late into the summer. An alternative explanation for the variation in SCPs we observed across habitats is the potential for differences in gut contents given the range of holding periods (12-72 hours, Table S1). Because gut contents can include efficient ice nucleating agents (and raise SCPs; Danks, 2008), we would expect higher SCPs for populations that were held for the shortest amount of time if this was confounding our results. However, Tetonica Pond had one of the shortest holding periods (12 hours) yet the lowest SCP (Tables 1, S1).

Because we tested SCPs in the absence of ice, it is likely that the inoculative freezing temperatures of our focal species are much higher than the values we report. For example, when in contact with ice, the SCP of *N. arctica* is -1.5 °C, but in its absence, *N. arctica* supercools to - 7.8 °C (Walters et al. 2009), a value that aligns with the range we observed (−7.5 to -5.2 °C; Table 1). Our ice enclosure experiment results also support the potential for higher inoculative freezing temperatures for *L. tumana* nymphs in contact with ice. Indeed, nymphs collected in late August did not survive being enclosed in ice, even at temperatures just below freezing (−0.5 °C). It is unclear, however, if the nymphs did not survive due to inoculative freezing or something else (e.g., internal mechanical damage or hypoxia). Nevertheless, a similar result was found for *N. arctica*. In mid-August, less than 20% of *N. arctica* nymphs survived a two-hour exposure to - 1.5°C. Yet, by late September when stream temperatures are colder and closer to 0°C, more than 80% survived a 7-day exposure to -6.2 °C, suggesting the presence of plasticity in cold tolerance (Walters et al. 2009). Although testing for seasonal plasticity in freeze tolerance was beyond the scope of our study, when typical summer conditions prevail (e.g., T_MEAN_ > 5°C), *L. tumana* nymphs from Lunch Creek are intolerant of even mild subzero temperatures. But unlike the Chandalar River in Alaska, where *N. arctica* resides and temperatures reach -10 °C at the sediment-water interface in winter, streams in the high Rocky Mountains appear to rarely, if ever, fall below 0 °C. Indeed, for the streams in this study for which we have full year thermographs (Figure S2), only Lunch Creek approached 0 °C at any point during the year. This thermal buffering likely reflects high levels of insulating snow in the region. However, our reported temperatures reflect the thermal conditions of the main stream channel. At stream margins, where flows are slower and less water is available for thermal buffering, temperatures may be quite different.

In this study, we showed that SCPs vary among species, populations, and habitat types for stonefly nymphs inhabiting Rocky Mountain headwaters. The population most likely to experience winter freezing stress also exhibited the lowest SCP, suggesting a potential ecological role for SCP in freeze avoidance among alpine stoneflies. Our results, of course, apply only to nymphs, one of three life stages in stoneflies: eggs, nymphs (i.e., larvae), and adults. We focused on nymphs because this stage is the key period when most growth occurs. High-elevation nemourid stoneflies appear to spend at least one winter as larvae (Figure S1), and statistical models indicate that species’ vulnerability to climate change can be greatly underestimated when the larval period is overlooked (Levy et al. 2015). With evidence across stonefly species for cryoprotective dehydration greatly lowering the freezing point of eggs (Gehrken & Sømme 1987), the ability for nymphs to survive being enclosed in ice (Walters et al. 2009), and adults that emerge in winter (e.g., Bouchard Jr. et al. 2009), it is clear that stoneflies can survive freezing conditions across all life stages. However, the degree to which freeze tolerance is common or rare, and how cold tolerance varies across life stages within species are key questions for future study.

As climate change proceeds, glaciers and perennial snowfields are receding, driving reduced mountain streamflow (Hotaling et al. 2017) and habitat reductions for high-elevation aquatic invertebrates (Domisch et al. 2011, Muhlfeld et al. 2020). Loss of meltwater, higher ambient temperatures, and rapid contemporary stream warming (e.g., Niedrist & Füreder 2020) are raising the risks for headwater biodiversity, including many stoneflies that are considered cold-water stenotherms (de Figueroa et al. 2010, Domisch et al. 2011, but see Hotaling et al. 2020). Reduced streamflow will lower the thermal buffering capacity of streamwater (Shah et al. 2020) and, despite warmer winter temperatures, it will also increase the potential for freezing stress (Williams et al. 2015). This effect will likely be compounded by decreasing levels of insulating snowpack as more winter precipitation falls as rain instead of snow (e.g., Huntington et al. 2004). Specifically, if snowpack accumulates later or melts off earlier, freezing risk in transitional periods between seasons (e.g., November and June in the Northern Hemisphere) may increase. Thus, a greater risk of freezing may represent an overlooked climate change threat to alpine aquatic biodiversity. Although only two stoneflies are known to actively mitigate freezing stress during non-adult stages (Gehrken 1989, Walters et al. 2009), it is unclear if this reflects general stonefly biology or a lack of investigation.

## ACKNOWLEDGEMENTS

We acknowledge long-term funding from the University of Wyoming-National Park Service Research Station (http://uwnps.org). S.H. and J.L.K were supported by NSF awards OPP-1906015 and IOS-1557795. A.A.S. was supported by an NSF Postdoctoral Research Fellowship in Biology (DBI-1807694). M.E.D. was supported by NSF awards DEB-1457659, OIA-1826834, and EF-1921562. H.A.W. was supported by a grant from the Montana Water Center. We thank Taylor Price and Lydia Zeglin for field support. Jordan Boersma, Georg Niedrist, and Brent Sinclair provided valuable comments on the manuscript.

