## Supplementary Materials for "Cold tolerance of mountain stoneflies (Plecoptera: Nemouridae) from the high Rocky Mountains"

**Figures:**

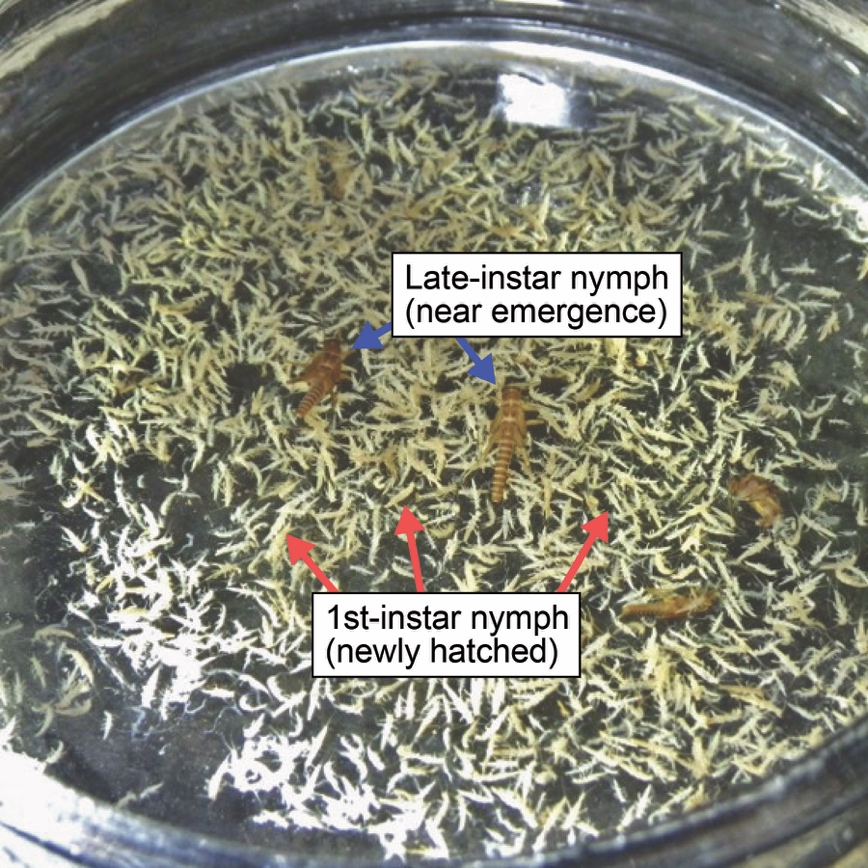

**Figure S1.** Two size classes of *Lednia tumana* present in a quantitative September sample of the macroinvertebrate community from Lunch Creek, Montana, USA.

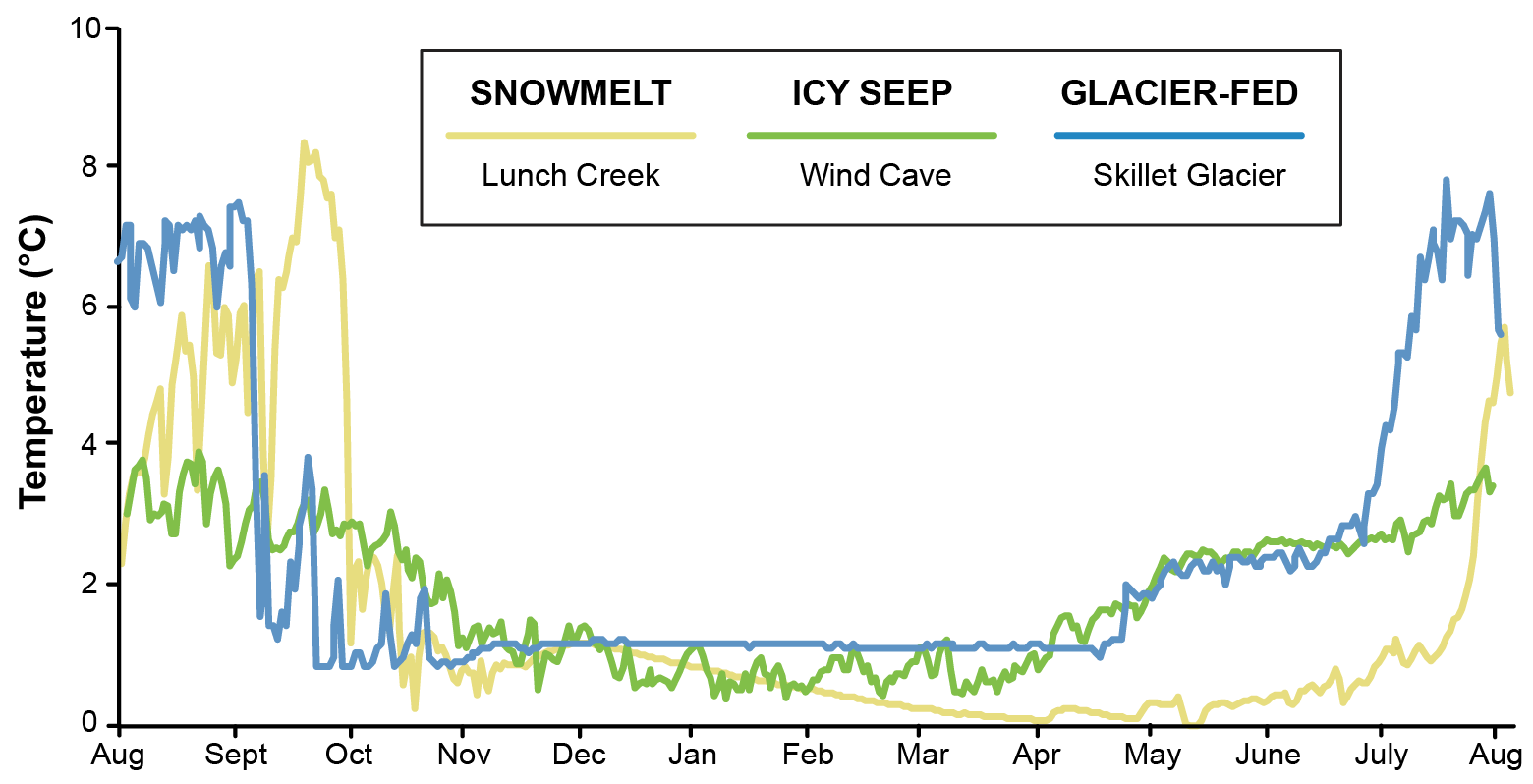

**Figure S2.** Daily mean temperatures for the three streams included in this study with a complete year of temperature data. Wind Cave and Skillet Glacier data are from 2016-2017. Lunch Creek was measured in 2013-2014.

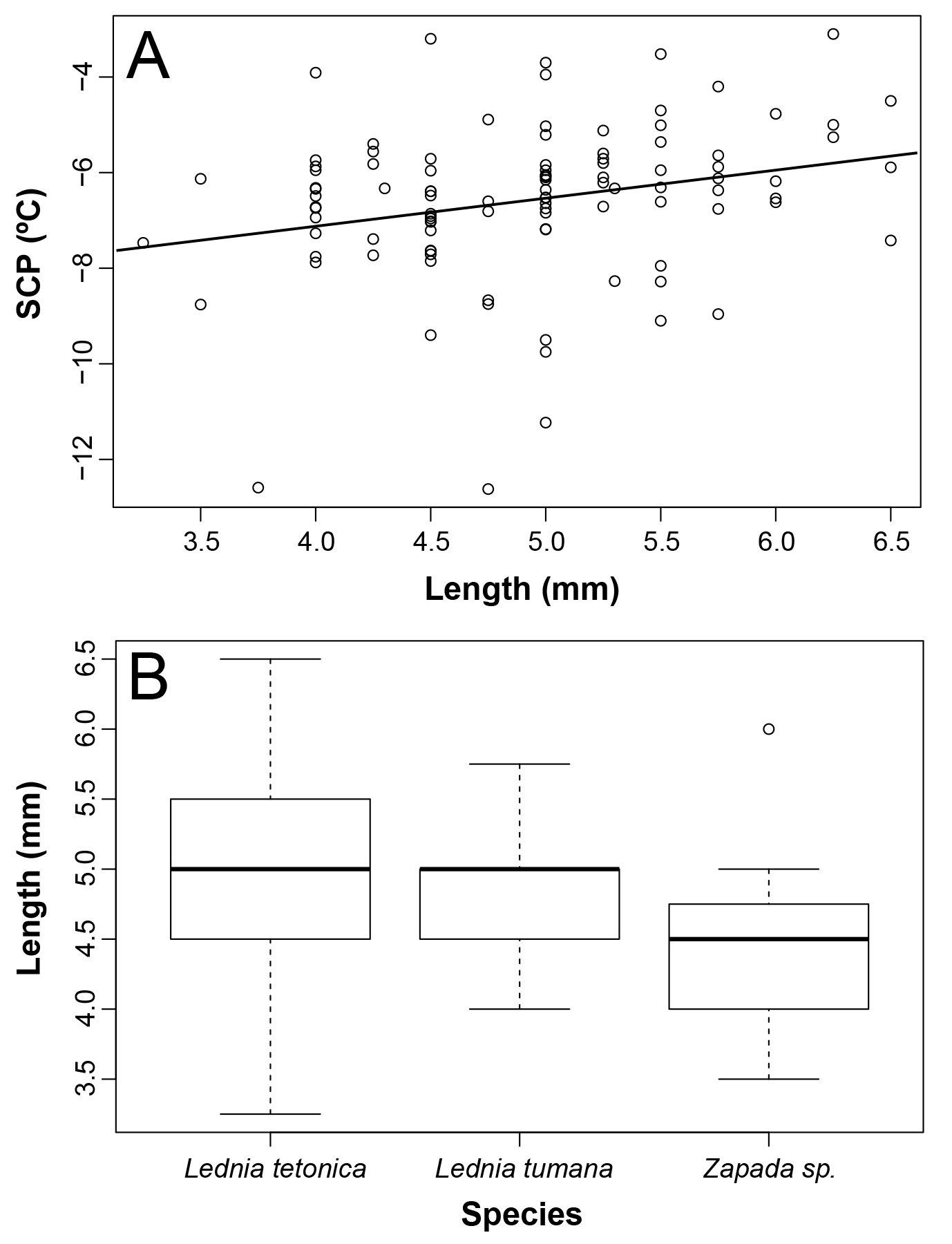

**Figure S3.** (A) Stonefly nymph length versus super cooling point (SCP). Longer nymphs had higher SCPs (*P*, ANOVA = 0.009). (B) Boxplots of body length by species included in the study.

**Tables:**

**Table S1.** Populations and species included in this study. Holding period = time (hours) that specimens were held at 3°C with no access to food before testing. *N* = sample size for each population. Ice-free = the approximate day in summer 2018 when the site became free of seasonal snow as measured with European Space Agency Sentinel-2 data (Copernicus Sentinel 2, 2018). Elevations are in meters (m) and lengths are in millimeters (mm).

| Population | Lat., Long. | Elevation | Ice-free | Length | Holding period | *N* |
| --- | --- | --- | --- | --- | --- | --- |
| Lunch Creek | 48.705, -113.704 | 2101 | 12 July | 4.9 ± 0.5 | 72 | 25 |
| Wind Cave | 43.667, -110.956 | 2606 | 12 June | 4.4 ± 0.6 | 48 | 20 |
| Mount Saint John | 43.792, -110.778 | 2686 | 5 July | 5.6 ± 0.7 | 12 | 19 |
| Cloudveil Dome | 43.725, -110.796 | 2928 | 1 Aug | 4.5 ± 0.5 | 12 | 18 |
| Skillet Glacier | 43.839, -110.757 | 2768 | 20 July | 4.6 ± 0.6 | 12 | 12 |
| Tetonica Pond | 43.788, -110.795 | 2804 | 15 July | 4.9 ± 0.5 | 12 | 17 |

**Table S2.** Tukey-adjusted differences in supercooling point for pairs of populations. Only significant differences (*P* < 0.05) are shown. See Table 1 for full population names.

| Population pair | Difference | Lower | Upper | *P*-adjusted |
| --- | --- | --- | --- | --- |
| MSJ-TP | 1.76 | 0.28 | 3.23 | 0.009 |
| SG-TP | 2.33 | 0.67 | 4.00 | 0.001 |
| MSJ-WC | 1.59 | 0.14 | 3.04 | 0.022 |
| SG-WC | 2.16 | 0.52 | 3.81 | 0.003 |
| SG-CD | 1.80 | 0.14 | 3.47 | 0.024 |
